# Normalizing automatic spinal cord cross-sectional area measures

**DOI:** 10.1101/2021.09.30.462636

**Authors:** S. Bédard, J. Cohen-Adad

## Abstract

Spinal cord cross-sectional area (CSA) is a relevant biomarker to assess spinal cord atrophy in various neurodegenerative diseases. However, the considerable inter-subject variability among healthy participants currently limits its usage. Previous studies explored factors contributing to the variability, yet the normalization models were based on a relatively limited number of participants (typically < 300 participants), required manual intervention, and were not implemented in an open-access comprehensive analysis pipeline. Another limitation is related to the imprecise prediction of the spinal levels when using vertebral levels as a reference; a question never addressed before in the search for a normalization method. In this study we implemented a method to measure CSA automatically from a spatial reference based on the central nervous system (the pontomedullary junction, PMJ), we investigated various factors to explain variability, and we developed normalization strategies on a large cohort (N=804).

Cervical spinal cord CSA was computed on T1w MRI scans for 804 participants from the UK Biobank database. In addition to computing cross-sectional at the C2-C3 vertebral disc, it was also measured at 64 mm caudal from the PMJ. The effect of various biological, demographic and anatomical factors was explored by computing *Pearson’s* correlation coefficients. A stepwise linear regression found significant predictors; the coefficients of the best fit model were used to normalize CSA.

The correlation between CSA measured at C2-C3 and using the PMJ was *y* = 0.98*x* + 1.78 (*R*^*2*^ = 0.97). The best normalization model included thalamus volume, brain volume, sex and interaction between brain volume and sex. With this model, the coefficient of variation went down from 10.09% (without normalization) to 8.59%, a reduction of 14.85%.

In this study we identified factors explaining inter-subject variability of spinal cord CSA over a large cohort of participants, and developed a normalization model to reduce the variability. We implemented an approach, based on the PMJ, to measure CSA to overcome limitations associated with the vertebral reference. This approach warrants further validation, especially in longitudinal cohorts. The PMJ-based method and normalization models are readily available in the Spinal Cord Toolbox.

## 1. Introduction

Various neurodegenerative diseases such as multiple sclerosis (MS) are associated with spinal cord (SC) atrophy, which is caused by demyelination, neuronal and/or axonal loss (Bonacchi et al., 2020; Lukas et al., 2013). New techniques have now become available through recent advancement in magnetic resonance imaging (MRI) and are relevant to assess SC atrophy (Moccia et al., 2019).

SC atrophy at the upper cervical levels can be defined within its cross-sectional area (CSA) (Lukas et al., 2013). The use of this metric is yet still limited due to considerable inter-subject variability. Finding factors that contribute to the observed variability is crucial to improve sensitivity and specificity of SC CSA and to develop normalization strategies.

Various studies have explored the correlation between SC CSA and demographic, anatomical and biological factors. Sex was a relevant factor to explain SC CSA variability, with females having significantly smaller SC CSA than males (Engl et al., 2013; Nico Papinutto et al., 2020; Solstrand Dahlberg et al., 2020). While the majority of studies have reported this significant effect, Fradet et al. (2014) found it to be an irrelevant factor and Papinutto et al. (2015) did observe the trend, but no statistical difference was found. However, the absence of a statistical difference could be explained by the relatively small sample size (30 participants).

Regarding the effect of age, a decrease of SC CSA was previously reported (Engl et al., 2013; Ishikawa et al., 2003; Kato et al., 2012; Nico Papinutto et al., 2015, 2020). However, the effect was not significant (Engl et al., 2013; Nico Papinutto et al., 2015, 2020). This trend is accentuated for older populations, but the effect of age is still small (Engl et al., 2013). An increase of SC CSA values followed by a decrease at 45 years old was also reported, but the effect was not significant (Nico Papinutto et al., 2020). The effect of age on SC CSA needs further investigation, small sample size and narrow range of age are limiting factors to assess the effect of age on SC CSA.

As for height and body weight, no significant effect on SC CSA was found according to recent studies (Nico Papinutto et al., 2015, 2020; Solstrand Dahlberg et al., 2020). The effect of height may be driven by sex differences (Nico Papinutto et al., 2020). Body mass index (BMI) was also tested as a normalization strategy for SC volume, but results were inconclusive as inter-subject variability was increased (Sanfilipo et al., 2004). In another study, no correlation was found with BMI by Solstrand Dahlberg et al.(2020).

Strong correlation between brain metrics and SC CSA were reported in prior studies (Engl et al., 2013; Nico Papinutto et al., 2015, 2020; Solstrand Dahlberg et al., 2020). White matter (WM) volume significantly explains upper cervical area variability as opposed to cerebrospinal fluid (CSF) volume, which was not significant according to Engl et al. (2013). In addition, brain volume correlated strongly with SC CSA as for intracranial volume (Nico Papinutto et al., 2015; Solstrand Dahlberg et al., 2020). Papinutto et al. (2020) also found this effect with V-scale (scaling factor for head size normalization) and considered it as the most promising factor for normalization strategies. Intracranial volume was also considered for normalization of SC volume but had limited utility since it generally diminished the ability to detect clinical-radiological correlations (Healy et al., 2012; Oh et al., 2014). Since SC CSA is a useful metric to assess SC atrophy and brain volume changes have also been associated with those pathologies, intracranial volume would be a better factor to consider for normalization strategies (Kesenheimer et al., 2021). A strong correlation was also found between thalamus volume and SC CSA by Solstrand Dahlberg et al. (2020). Axial canal area was also a significant factor and promising for normalization strategies, but has not been explored yet by many (Kesenheimer et al., 2021; Nico Papinutto et al., 2020). A notable difficulty for computing the axial canal area is the ability to properly segment it.

Only a few of the previous cited works have explored normalization strategies for SC CSA. SC length was a relevant factor for SC volume normalization compared to intracranial volume (Healy et al., 2012; Oh et al., 2014). Mean SC volume in healthy participants was also used to normalize SC volume of patients with MS (Ruggieri et al., 2021). Regarding SC CSA, age, intracranial volume and sagittal vertebral area were the most promising independent variables found by Papinutto et al. (2015), but the method was based on 30 participants only. Another model later including V-scale, axial canal product (product of maximum axial anterior-posterior and lateral diameters of the cervical SC) based on 129 participants significantly reduced SC CSA variability (Nico Papinutto et al., 2020). Brain WM volume, sex and spinal canal area formed a relevant normalization strategie, total intracranial volume could also replace brain WM volume for subjet with diseases that affect WM. The main limitation here was the relatively small number of participants (N=61) (Kesenheimer et al., 2021). As we can observe, brain/skull metrics and sex are important factors to consider in a possible normalization method. Also, the mentioned normalization methods are only reported in the related papers; they are not easily reusable to integrate directly within analysis pipelines.

In addition to the biological-derived normalization strategy, previous studies have reported a variability in CSA measures associated with the MRI acquisition parameters, and segmentation method (Cohen-Adad et al., 2021; Kearney et al., 2014; Nico Papinutto & Henry, 2019).

The majority of the studies regarding SC CSA use vertebral levels as an anatomical reference. However, the prediction of the spinal segments is imprecise since it is based on vertebral levels, adding variability (Cadotte et al., 2015). Inferring neuroanatomic positions with vertebral bodies doesn’t consider neck flexion and extension. Segmental nerve rootlets would provide a proper identification of the spinal segments, it is however difficult to identify and requires high resolution T2w scans and an expert rater to identify them. A few studies have attempted to bypass the vertebral-based limitation using the distance from the pontomedullary junction (PMJ) (Cadotte et al., 2015; Stroman et al., 2008).

While SC CSA variability across participants was shown to be associated with multiple demographic, anatomical and biological factors (Kesenheimer et al., 2021; Nico Papinutto et al., 2015, 2020; Solstrand Dahlberg et al., 2020), no previous studies have addressed the variability associated with limitation of vertebral based SC CSA measurement in the search for a normalization method.

In this study we quantify the contribution of various factors on the inter-subject variability in cervical SC CSA measurements. We notably introduce a method to replace the traditional vertebral-based referencial system by an anatomical reference from the central nervous system. More precisely, we (1) establish an automatic MRI data processing pipeline to compute SC CSA, (2) process MRI data from a subset (N=804) of the UK Biobank database, (3) introduce a method to automatically detect the PMJ and use it as a referential system to measure SC CSA, (4) develop a statistical model and normalization method for SC CSA measurements, (5) make this model readily available to use via the open-source Spinal Cord Toolbox (SCT) software (De Leener et al., 2017).

## 2. Material and methods

### 2.1. Demography

1,000 participants (48 to 80 years old, 56.3% female) were selected from the UK Biobank database. Not knowing the effect size we were after, we could not base this number on any reliable power analysis. Hence, the number of participants was selected as a compromise between the statistical power we wanted to achieve in comparison with the previously-published studies addressing similar scientific questions (typically < 300 participants) and the time required to manually validate each step of the processing pipeline (visual inspection, manual correction of SC segmentation and/or PMJ labeling and/or vertebral labeling).

Participants with a history of neurological diseases were excluded from the study. Fields from UK Biobank dataset included in the category *Nervous system disorders*^*1*^ were used to identify these participants. This brought the number of participants from 1,000 to 972.

### 2.2. Image acquisition

Data used for this study were unprocessed NIfTI T1w structural scans from the UK Biobank Brain Imaging dataset (Miller et al., 2016). Images were acquired in four different assessment centers on a Siemens Skyra 3T running VD13A SP4 with a standard Siemens 32-channel RF receive head coil. T1w structural scan has a field of view (FOV) of 208×256×256 with an isometric resolution of 1 mm^3^. The superior-inferior field of view of 256 mm typically covers down to C3 vertebral level, which is relevant for the present study as SC CSA was measured around the C2-C3 vertebral level. The UK Biobank data includes preprocessed data (corrected for gradient non-linearity and masked), however we could not use these data because the SC was masked out. We therefore used the unprocessed T1w images as input of the processing pipeline described in the next section.

### 2.3. Data processing

Data processing pipeline is based on spine-generic v2.6^2^ (Cohen-Adad et al., 2021) pipeline and SCT v5.4^3^ (De Leener et al., 2017). The processing pipeline is available on GitHub^4,5^ and is fully documented^6^.

Figure 1. presents an overview of the processing pipeline. First, all images were reoriented to right-inferior-posterior orientation. Since SC CSA is computed using T1w brain images and SC is in the periphery of the images’ FOV, gradient non-linearity distortions have a considerable effect on CSA measures, depending on participant positioning (N. Papinutto et al., 2018). Therefore, correction for gradient non-linearities was applied using *gradunwrap* from the HCP project (Glasser et al., 2013) and the coefficient file for the Siemens Skyra 3T gradient system. Then, the SC was segmented automatically using deep learning models with SCT’s `sct_deepseg_sc` (Gros et al., 2019). SC CSA was computed and averaged using SCT’s `sct_process_segmentation` with two different methods further explained below.

**Figure 1.**
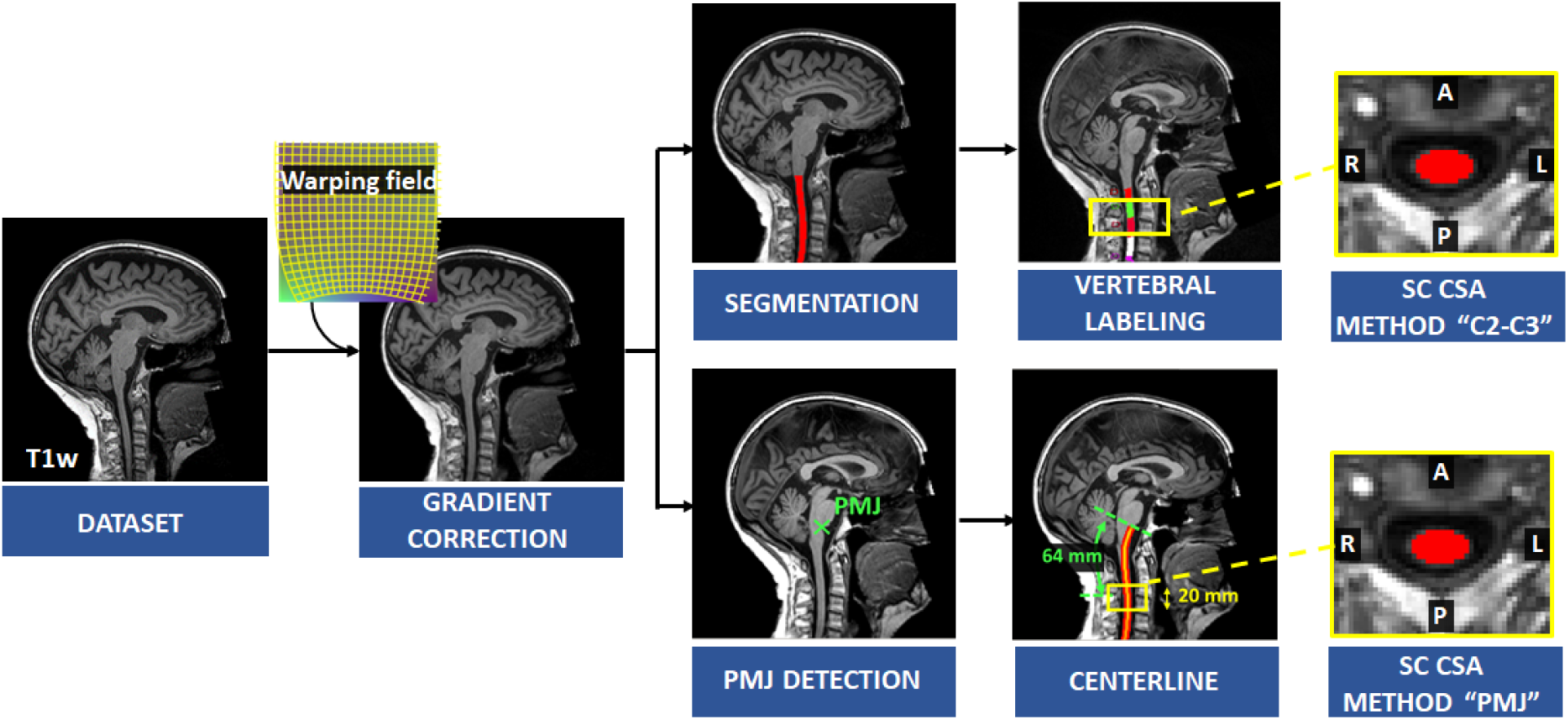
Overview of the processing pipeline. For each participant, correction for gradient non-linearities was applied on the T1w image and SC was segmented automatically. Vertebral levels were identified and SC CSA was computed at C2-C3 levels. PMJ was labeled followed by extracting the centerline to compute SC CSA at 64 mm from PMJ averaged on a 20 mm extent.

#### 2.3.1. CSA based on distance from neurological reference: pontomedullary junction

To overcome the limitation of SC segment prediction with vertebral bodies, SC CSA was measured from a distance of a neurological reference; we chose the PMJ (Cadotte et al., 2015; Stroman et al., 2008).

First, the PMJ was identified using SCT’S `sct_detect_pmj.` Briefly, a 2D support vector machine trained with histogram of oriented gradient features (HOG+SVM 2D classifier) was run on the mid-sagittal slice to detect the PMJ (Gros et al., 2017). Since the mid-sagittal slice does not necessarily correspond to the anatomical medial plane, with a sliding window centered on the first estimated PMJ coordinate, cross-correlation was computed within the window and its mirror image in the right-left orientation. We assumed that the maximum cross-correlation corresponds to the right-left symmetry slice. The HOG+SVM 2D classifier was run again on the updated medial plane.

Since the SC curvature associated with cervical lordosis varies across individuals, the distance from the PMJ was computed along the SC centerline following the arc-length. SC segmentation normally doesn’t go as high as the PMJ. The PMJ coordinate was then added to the SC segmentation prior to extracting the SC centerline. Linear interpolation and smoothing were used to extract the SC centerline. SC CSA was computed at 64 mm from PMJ along the centerline slice-wise, corrected for angulation and then averaged on a 20 mm extent as presented in **Figure 2**. The 64 mm value corresponds to the mean distance between C2-C3 disc and PMJ.

**Figure 2.**
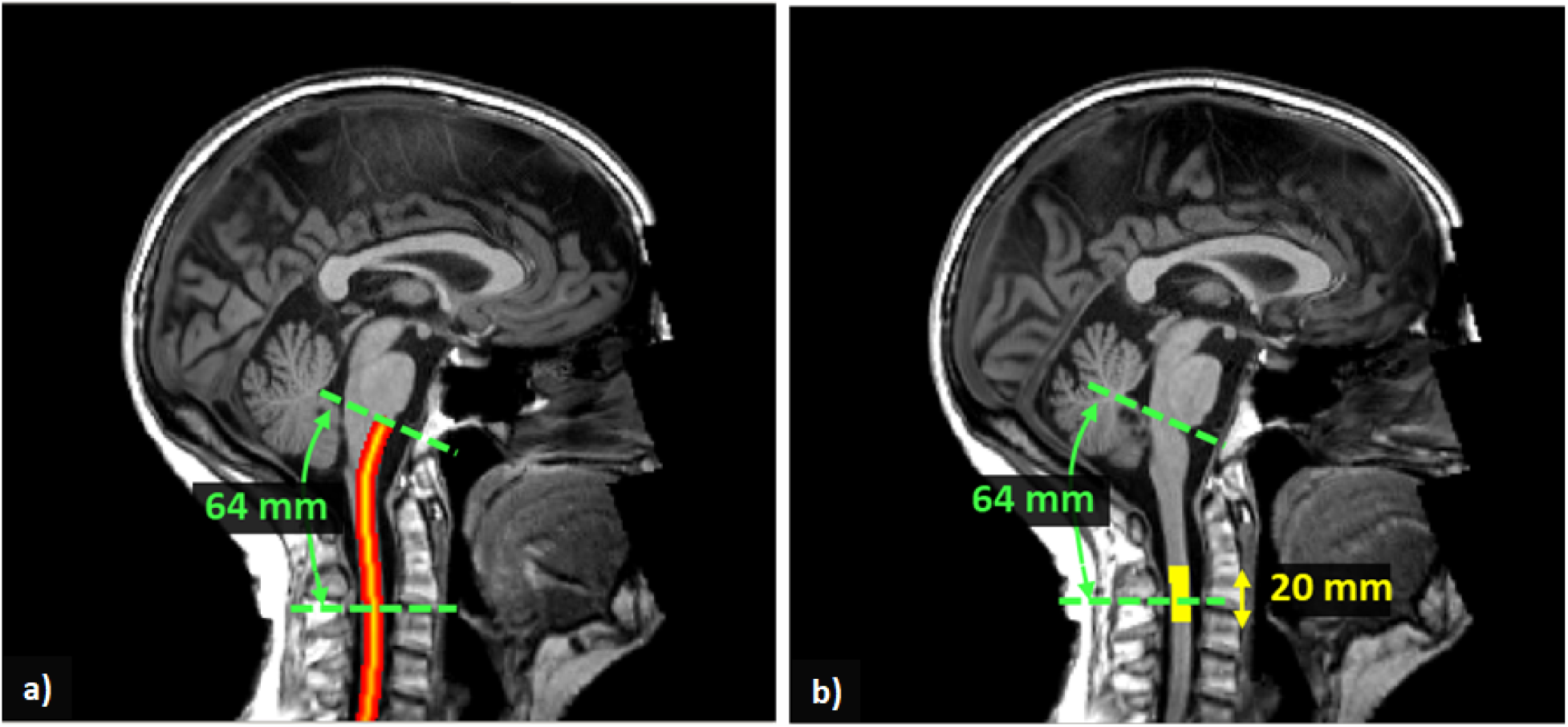
**a)** SC centerline from PMJ. SC centerline was extracted from SC segmentation and PMJ label using linear interpolation and smoothing. The distance from PMJ is measured along the centerline following the arc-length. **b)** Extent mask to average SC CSA. At 64 mm from PMJ, CSA was computed slice-wise, corrected for angulation and averaged within a 20 mm extent. The extent mask is centered at 64 mm from PMJ.

#### 2.3.2. CSA based on C2-C3 vertebral levels

The second and more popular method to compute the SC CSA was to use C2-C3 vertebral levels as an anatomical reference for spinal segments. Vertebral levels were identified using `sct_label_vertebrae` (Ullmann et al., 2014) and then averaged slice-wise, corrected for angulation between the SC and the slice and then averaged at C2-C3 vertebral levels.

### 2.4. Quality control

All SC segmentations, vertebral and PMJ labels were inspected and validated following a procedure^7^ inspired by the spine-generic project^8^ (Cohen-Adad et al., 2021). According to the quality of the segmentations and labels, manual corrections were applied. If data presented poor quality (ghosting, excessive motion, bad field of view placement), it was excluded from the statistical analysis. Examples of excluded images are listed in https://github.com/sct-pipeline/ukbiobank-spinalcord-csa/issues/61. This brought the number of participants from 972 to 826. After all manual corrections, we re-ran the analysis to obtain valid SC CSA values.

Processing was distributed across 40 CPU cores (one participant per CPU core) using `sct_run_batch` on a 64-core CPU cluster. Total processing time was 01h53m59s.

### 2.5. Statistical Analyses

#### 2.5.1. Comparison of CSA measure with both methods

To study the relationship between the PMJ-based and vertebral-based CSA measures, we built a scatterplot and derived a model. We also computed the distance between the C2-C3 disc and PMJ. Mean, standard deviation, median and coefficient of variation (COV) for both CSA measures were computed.

For the rest of the study, we used the PMJ-based CSA because it uses a coordinate system intrinsic to the central nervous system and thus represents a more faithful assignment/labeling of the spinal level under investigation.

#### 2.5.2. Correlations with physical and brain measures

In this study, the effect of sex, age, physical measures and brain measures on SC CSA was explored by calculating *Pearson’s* correlation coefficients. Physical measures included height and weight. Brain measures included brain WM volume, brain gray matter (GM) volume, brain volume, brain volume normalized for head size, thalamus volume and ventricular cerebrospinal fluid (CSF) volume. All data (images and demographic measures) were acquired at the same time for each participant and were available in the UK Biobank database.

#### 2.5.3. Effect of sex and age

To assess the effect of sex on SC CSA, a T-test was performed to establish if the mean CSA for male and female has a significant difference. To assess the effect of age on SC CSA, we fitted a linear and quadratic regression, *R*^*2*^ were reported.

#### 2.5.4. Multilinear regression

The effect of all candidate predictors was evaluated using a multilinear analysis. Sex was included as a dichotomous variable to the regression. Any participant missing a parameter was excluded from this analysis. This brought the number of participants from 826 to 804. To select the relevant predictors of the multilinear regression, a stepwise method was used. The predictors are added to the model from the highest correlation with CSA to the lowest if they are significant (*p-value* < 0.05). After each addition of predictors, the significance of the current parameters was computed again, and parameters with a *p-value* > 0.05 were excluded from the model (Toutenburg, 1969). The level of significance was the same for both entry and exit tests. To validate the model, we proceeded to a residual analysis and computed *R*^*2*^.

*Pearson’s* correlation coefficients between candidate predictors was used to choose which parameter to include in the stepwise model due to possible collinearity between parameters.

#### 2.5.5. Normalization method

With the best multilinear regression fit, a regression-based residual method (Nico Papinutto et al., 2020; Sanfilipo et al., 2004) was developed using the significant predictors, as described in the following equation:

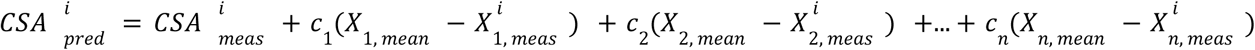

Where 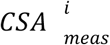 is the computed SC CSA value from a given participant *i*, 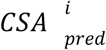 is the normalized CSA value, *c*_*j*_are the coefficients of the multilinear regression, *X*_*j,mean*_ are the mean values of all significant predictors, 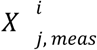 are values for the given participant’s predictors, for *j* predictors (Nico Papinutto et al., 2020).

Since this method assumes that the regression line slopes are parallel for both groups for sex (Sanfilipo et al., 2004), the interaction between signifcant predictors and sex was also explored afteward. If the interaction term was significant, it was added to the model. The interaction term corresponds to the predictor multiplided by sex (0 or 1) as we can see in the following equation:

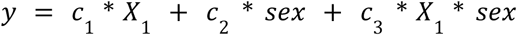

The effect of normalization was then evaluated by comparing the COV of the normalized SC 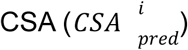 and the measured 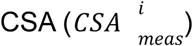.

## 3. Results

### 3.1. SC CSA

In this study we compared SC CSA measured using the PMJ as a reference or the C2-C3 disc as a reference (more popular). When using the PMJ as a reference (64 mm caudal to the PMJ), the CSA ranged between 51.9 and 95.6 mm^2^ (mean ± SD: 66.2 mm^2^ ± 6.69 mm^2^). The COV was 10.09%. When using C2-C3 vertebral levels as a reference, the CSA ranged between 51.5 and 96.9 mm^2^ (mean ± SD: 66.4 mm^2^ ±6.61. mm^2^). The COV was 9.96 %.

**Figure 3a)** shows the relationship between CSA at 64 mm from the PMJ and CSA at C2-C3 vertebral levels. The linear regression led to a *R*^*2*^ of 0.97. **Figure 3b)** shows a scatterplot of the distance from the PMJ and C2-C3 disc. Mean distance from the PMJ to the C2-C3 disc is 64.37 ± 5.53 mm.

**Figure 3.**
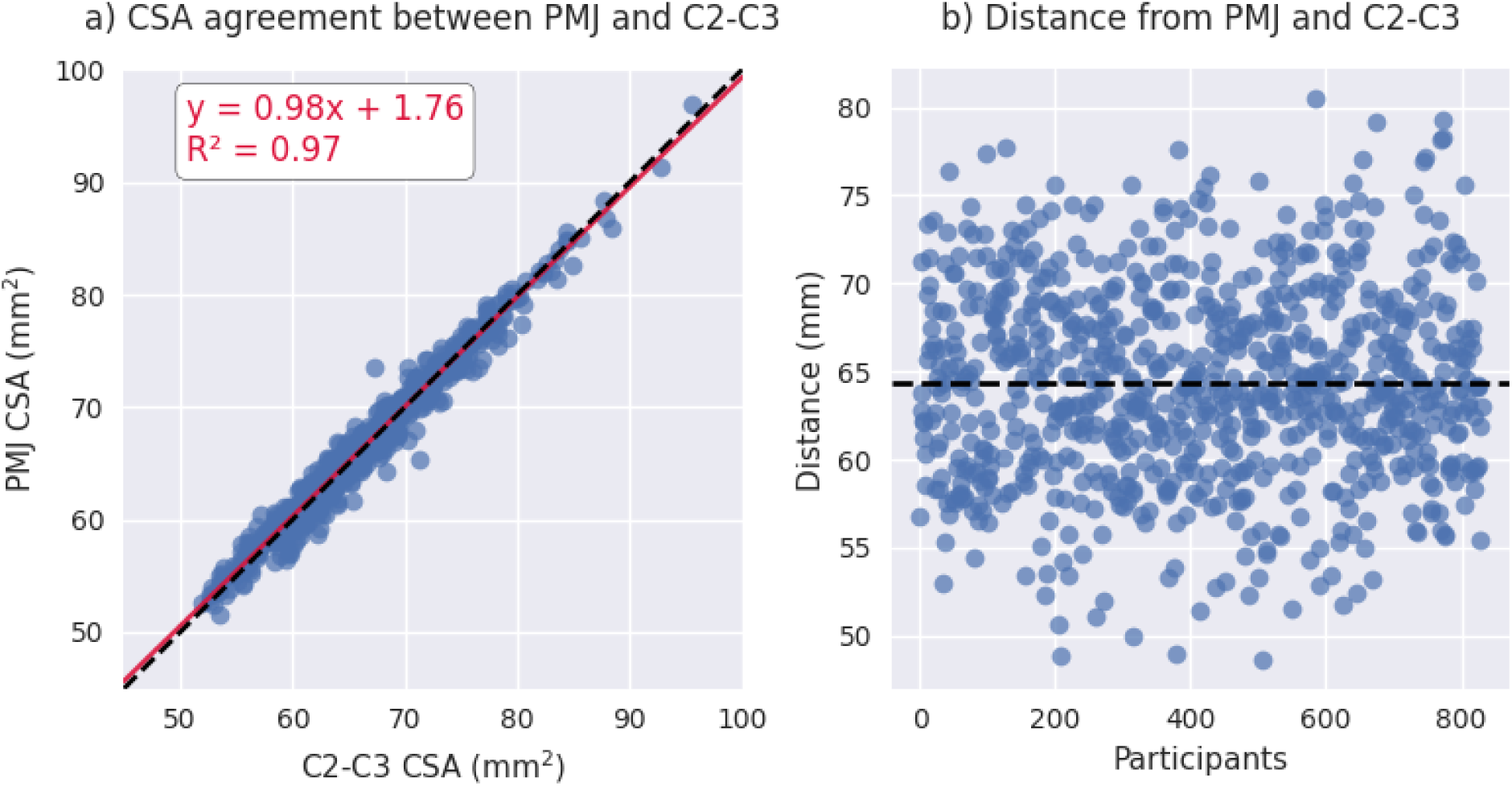
**a)** Scatterplot of PMJ-based CSA at 64 mm and vertebral-based CSA at C2-C3 vertebral levels. **b)** Scatterplot of the distance between the PMJ and the C2-C3 disc.

For the rest of the results, SC CSA will refer to PMJ-based CSA at 64 mm.

### 3.2. Statistical Analyses

#### 3.2.1. Correlations with physical and brain measures

We investigated the relationship of SC CSA with sex, age, physical and brain measures. Results of the correlation analysis (*Pearson’s*) are reported in **Table 1**. Note that this correlation matrix is not corrected for multiple comparisons because its purpose was only to explore existing correlations. In subsequent analysis (see *Multilinear regression*), a multivariate analysis will account for the number of regressors in the estimated *p-values*. Scatterplots of CSA and all parameters are shown in supplementary material (**S1**-**S8**). Ventricular CSF volume was the only parameter to present a non-significant correlation coefficient with SC CSA (*p-value* > 0.05). Thalamus, brain, brain WM and brain GM volume present the highests correlations out of all parameters (*Pearson’s r* > 0.4). Among parameters, we notice in particular a very strong correlation between thalamus volume and brain, brain WM and brain GM volume.

**Table 1.**
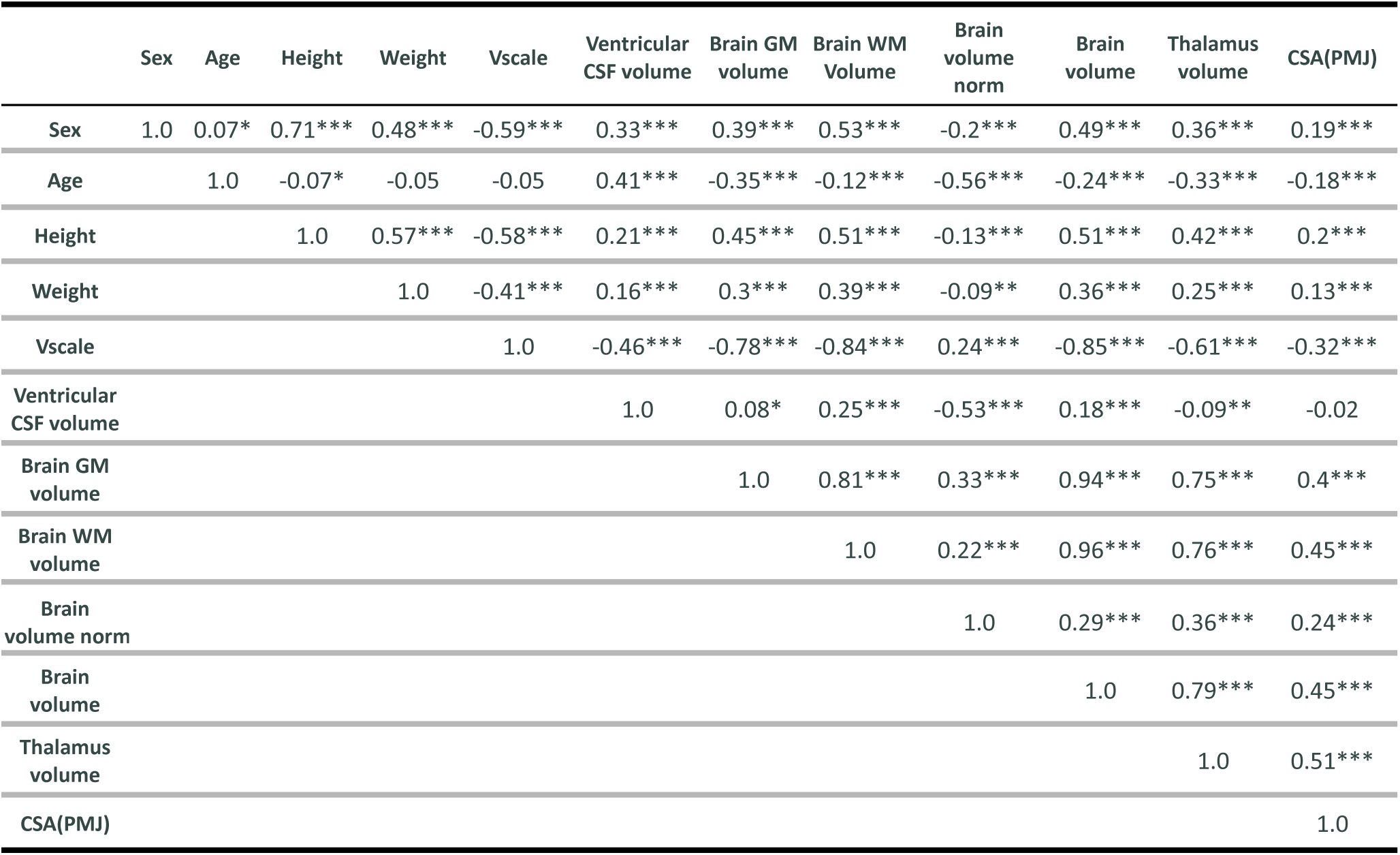
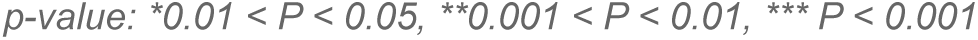
Pearson’s correlation among SC CSA, physical and brain measures.

|  | Sex | Age | Height | Weight | Vscale | Ventricular CSF volume | Brain GM volume | Brain WM Volume | Brain volume norm | Brain volume | Thalamus volume | CSA(PMJ) |
| --- | --- | --- | --- | --- | --- | --- | --- | --- | --- | --- | --- | --- |
| Sex | 1.0 | 0.07* | 0.71*** | 0.48*** | -0.59*** | 0.33*** | 0.39*** | 0.53*** | -0.2*** | 0.49*** | 0.36*** | 0.19*** |
| Age |  | 1.0 | -0.07* | -0.05 | -0.05 | 0.41*** | -0.35*** | -0.12*** | -0.56*** | -0.24*** | -0.33*** | -0.18*** |
| Height |  |  | 1.0 | 0.57*** | -0.58*** | 0.21*** | 0.45*** | 0.51*** | -0.13*** | 0.51*** | 0.42*** | 0.2*** |
| Weight |  |  |  | 1.0 | -0.41*** | 0.16*** | 0.3*** | 0.39*** | -0.09** | 0.36*** | 0.25*** | 0.13*** |
| Vscale |  |  |  |  | 1.0 | -0.46*** | -0.78*** | -0.84*** | 0.24*** | -0.85*** | -0.61*** | -0.32*** |
| Ventricular CSF volume |  |  |  |  |  | 1.0 | 0.08* | 0.25*** | -0.53*** | 0.18*** | -0.09** | -0.02 |
| Brain GM volume |  |  |  |  |  |  | 1.0 | 0.81*** | 0.33*** | 0.94*** | 0.75*** | 0.4*** |
| Brain WM volume |  |  |  |  |  |  |  | 1.0 | 0.22*** | 0.96*** | 0.76*** | 0.45*** |
| Brain volume norm |  |  |  |  |  |  |  |  | 1.0 | 0.29*** | 0.36*** | 0.24*** |
| Brain volume |  |  |  |  |  |  |  |  |  | 1.0 | 0.79*** | 0.45*** |
| Thalamus volume |  |  |  |  |  |  |  |  |  |  | 1.0 | 0.51*** |
| CSA(PMJ) |  |  |  |  |  |  |  |  |  |  |  | 1.0 |

*p*-value: \* $0.01 < P < 0.05$ , \*\* $0.001 < P < 0.01$ , \*\*\* $P < 0.001$

#### 3.2.2. Effect of sex and age

Participants in this study include 43.7 % of males and 56.3 % of females. **Figure 4** presents SC CSA violin plots for female and male with mean and standard deviation. We found a significant difference for CSA between female and male (*t* = - 5.37, *p-value* < 10^−7^).

**Figure 4.**
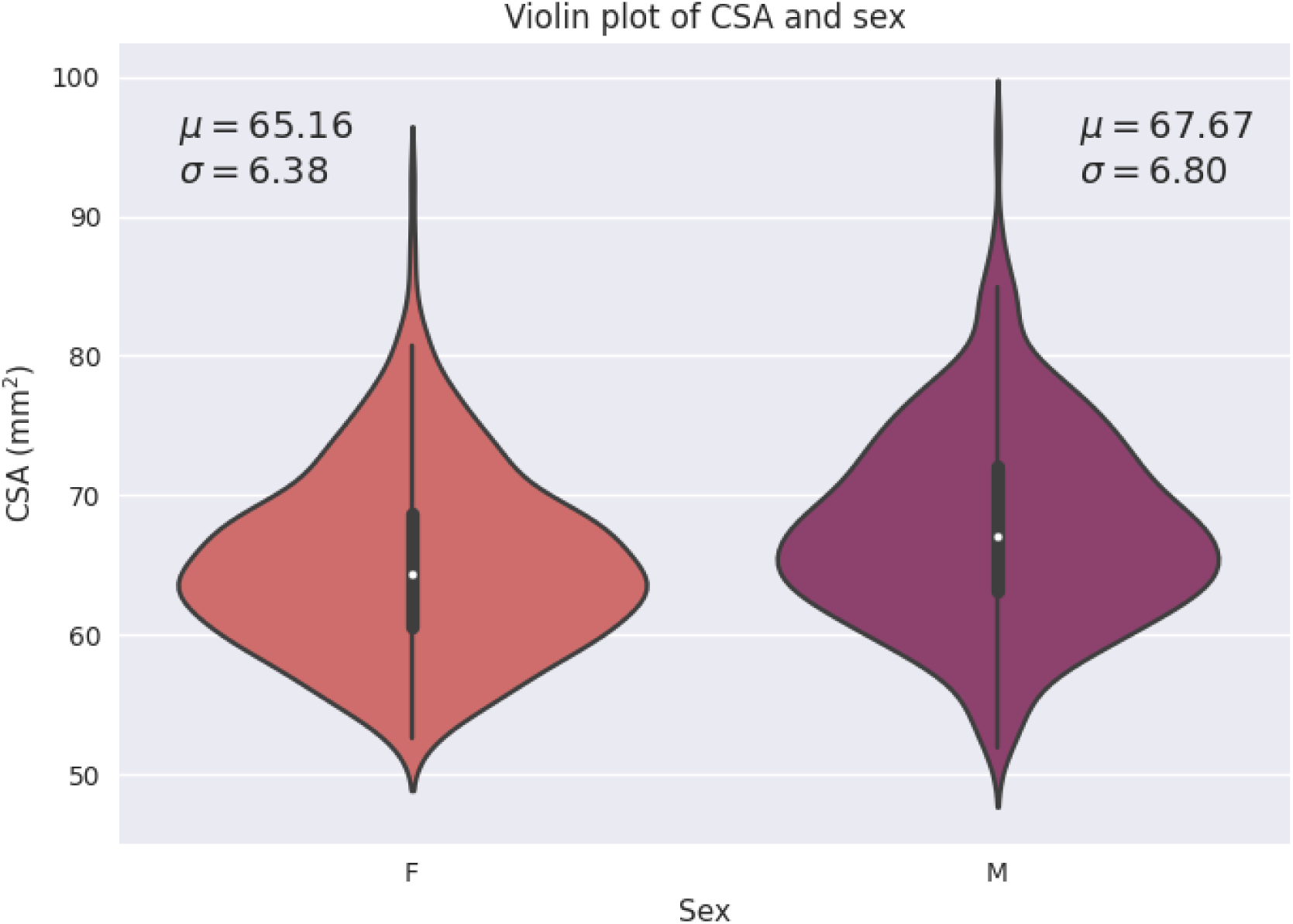
Violin plot of SC CSA for female and male with mean (µ) and standard deviation (σ) for each sex. F = female; M = male

To explore the relationship between age and SC CSA, we calculated a linear and quadratic fit. The age of the participants ranged between 48 and 80 years old. The linear fit is presented at **Figure 5**. The equation for the quadratic fit is:

**Figure 5.**
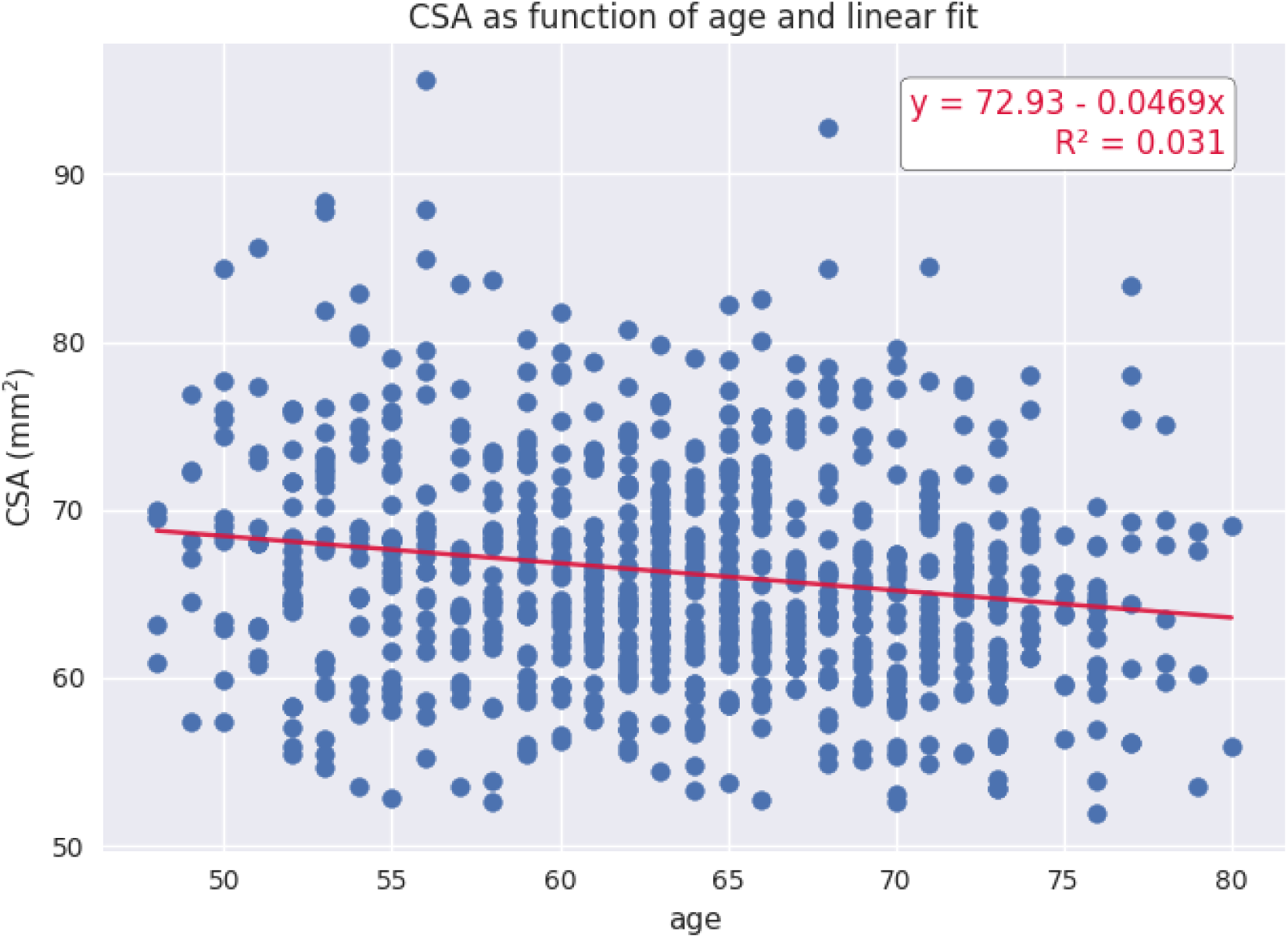
Linear fit for CSA as a function of age.

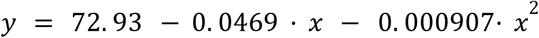

The constant and the linear coefficient are the same for both fits, and the quadratic coefficient is very small (9.07e-04): there is almost no quadratic trend.

#### 3.2.3. Multilinear regression

Based on the *Pearson’s* correlation analysis presented in **Table 1**, the following parameters were input in the stepwise linear regression: sex, height, weight, age, brain volume, ventricular CSF volume and thalamus volume. Since brain WM volume, GM volume and brain volume have a very strong correlation (0.94 and 0.96), brain WM volume and GM volume were not included in the model to avoid collinearity.

The stepwise method yielded a model including brain volume and thalamus volume. The resulting model is shown in **Table 2**. Adjusted *R*^*2*^ is 0.265. The model is significant (*p-value* < 0.0001).

**Table 2.**
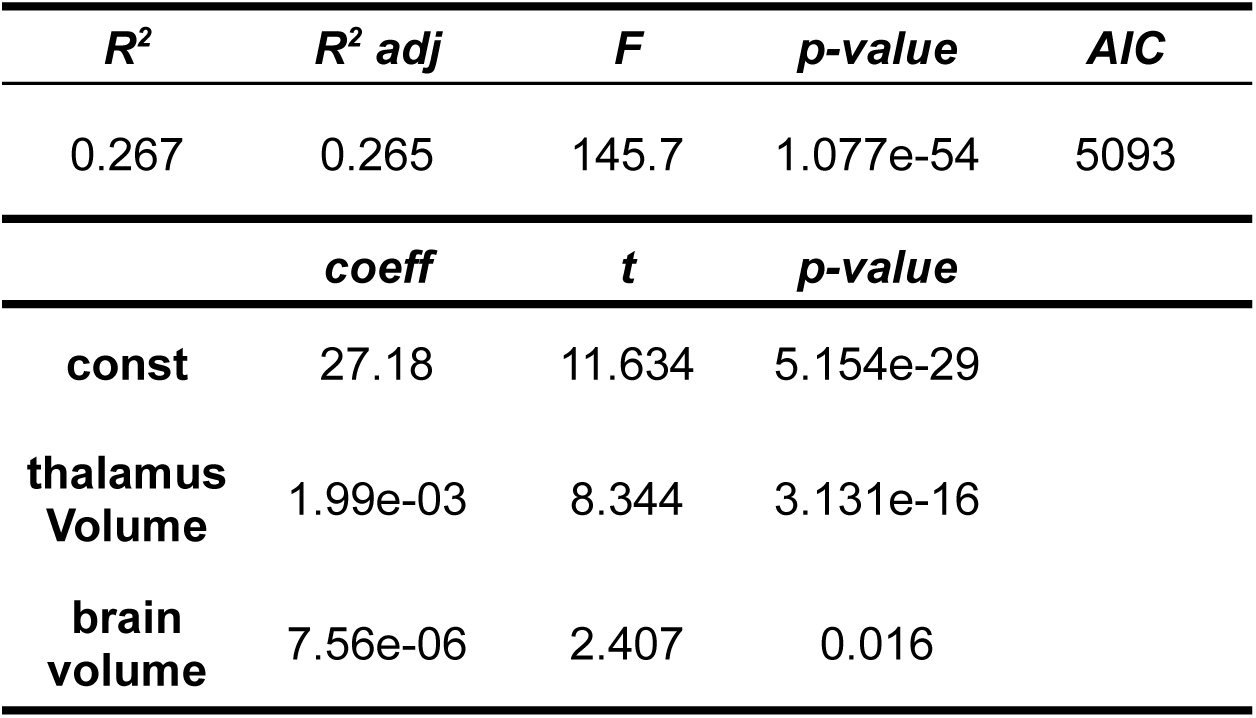
Multilinear Regression Analysis for SC CSA (N=804 participants).

### 3.3. Normalization

The presented model’s coefficients led to the following normalization equation with thalamus volume and brain volume and their respective means.

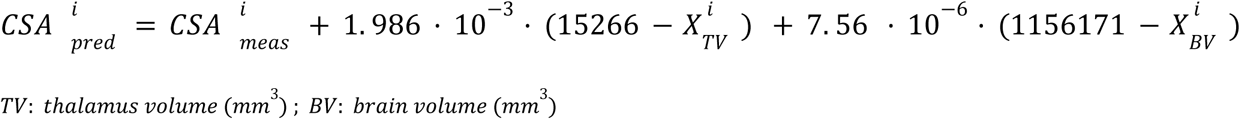

With the CSA normalization, COV went from 10.09 % 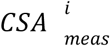 to 8.64 % 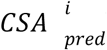, a reduction of 14.37%.

Figure 6. shows scatterplots of CSA with both predictors (brain volume and thalamus volume) separated for sex. Qualitatively, we observe that the slopes for female and male are different with the brain volume predictor, however the slopes are closer with the thalamus volume predictor. To quantitatively validate if the interaction coefficient is significant in the model, we computed the interaction of both predictors with sex. The interaction coefficient for brain volume was significant (*p-value* = 0.006) and sex was also significant when adding the interaction parameter (*p-value* = 0.005). For thalamus volume, the interaction coefficient was not significant (*p-value* = 0.227) neither was sex (*p-value* = 0.227). The interaction of brain volume and sex was therefore added to the previous model since it has a significant effect. The following equation presents the corresponding normalization equation:

**Figure 6.**
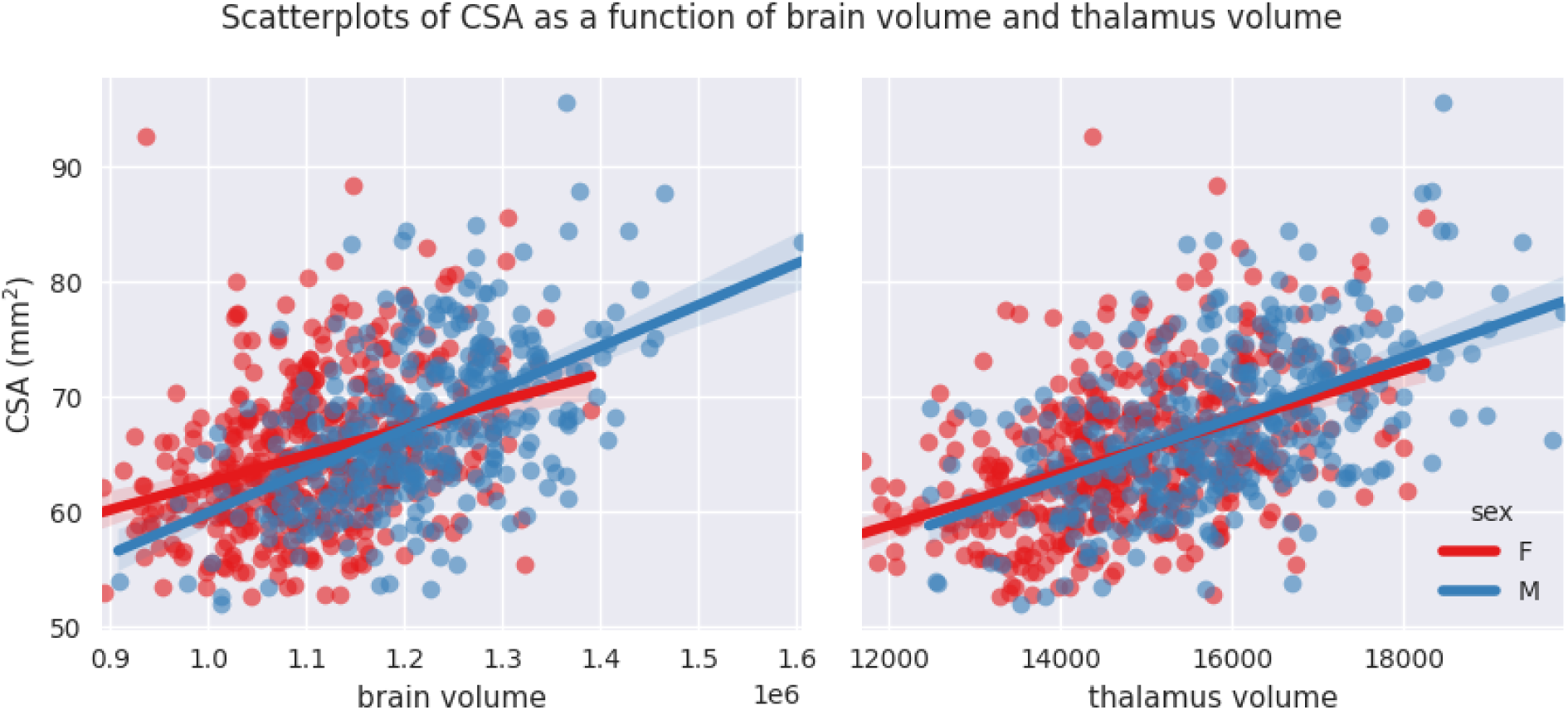
Scatterplots of CSA as a function of brain volume and thalamus volume separated for sex with linear fit.

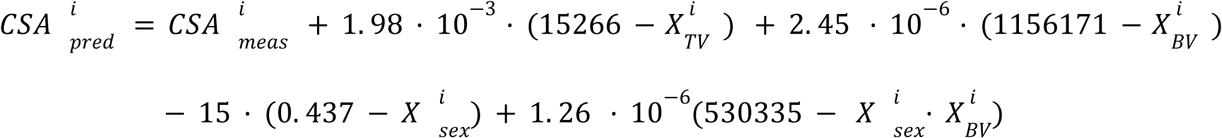

The model was significant (*p-value* < 0.0001). The COV of CSA went from 10.09% to 8.59%, a reduction of 14.85 %, a (modest) further reduction of COV compared to the model without sex interaction. Adjusted *R*^*2*^ was 0.271.

Since measuring thalamus volume is not always convenient, we also proposed a model without the thalamus volume as a predictor and kept only brain volume, sex and the interaction. The COV went from 10.09% to 8.96%, a reduction of 11.22%. Adjusted *R*^*2*^ was 0.209, the model was also significant (*p-value* < 0.0001). Even if this model is less performant than the one including the thalamus volume, we made it available for SCT users given that thalamus volume is not often measured in neuroimaging analysis pipelines. The normalization model has the following equation:

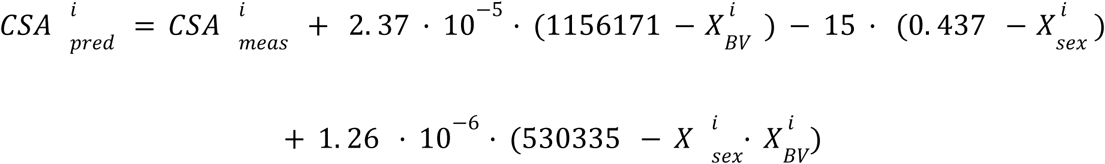

## 4. Discussion

In this work, we quantified the contribution of various factors on the inter-subject variability in cervical SC CSA measurements in 804 participants. We implemented a measurement method for SC CSA that uses the PMJ as opposed to the vertebral reference. Finally, we developed a normalization model which can reduce inter-subject variability by up to 14.85%. The method is available to use in the open-source software SCT (De Leener et al., 2017).

### 4.1. CSA results

We obtained mean CSA values of 66.2 mm^2^ and 66.4 mm^2^ for PMJ-based and vertebral-based CSA respectively, which is lower than what was reported in other studies (Kesenheimer et al., 2021; Nico Papinutto et al., 2020; Solstrand Dahlberg et al., 2020). Lower values could be explained by various factors. Firstly, the population studied here is relatively older than in other published studies (range: 48 to 80, mean: 64). Secondly, the segmentation method has an impact on defining the boundary between the SC and surrounding CSF. SCT’s `sct_deepseg_sc` is more conservative than other software in defining this border, which results in a smaller CSA (Lukas et al., 2021; Weeda et al., 2019). It is important to note that a systematic bias across software is not an issue when it comes to using CSA values for clinical studies: it only adds an offset and does not affect the precision of the measure. It is similar to a calibration problem. Thirdly, acquisition parameters (which drive image contrast) also influence CSA values (Cohen-Adad et al., 2021; Kearney et al., 2014; Nico Papinutto & Henry, 2019). Furthermore, gradient echo T1w acquisitions are prone to motion artifacts, which hamper the performance of SC segmentation. COV were 9.96% and 10.09% (C2-C3, PMJ), which is similar to what was observed in previous studies (Kesenheimer et al., 2021; Nico Papinutto et al., 2020; Solstrand Dahlberg et al., 2020).

### 4.2. PMJ-based CSA method

Regarding veterbral-based CSA and PMJ-based CSA, COVs are very similar (9.96% C2-C3, 10.09% PMJ) as for CSA values (see **Figure 3**). Therefore, there is no clear conclusion if the PMJ-based method is best for measuring CSA across individuals. However, for longitudinal studies (ie: intra-subject COV), given that head tilting might change across sessions, it is possible that a PMJ-based method is preferred. Subsequent study is needed to validate the relevance of a PMJ-based method for longitudinal studies.

Some limitations are associated with the PMJ-based CSA method. The use of the PMJ label to interpolate with the centerline is not the exact extrapolation of the centerline. SC curvature at the PMJ varies across individuals, which adds variability to the computed distance. PMJ label positioning across participants may also differ, also affecting the measured distance.

Moreover, the distance from PMJ doesn’t consider the fact that the SC length varies across individuals. Results from this study do not show a difference between SC CSA variability using a vertebral-based vs PMJ-based reference. A comparison with the nerve rootlets is necessary to assess which method ensures proper prediction of the spinal segments; it will be the subject of further investigations.

### 4.3. Correlation with SC CSA

Our analysis shows a strong correlation between brain volume, brain WM volume, brain GM volume and thalamus volume with CSA, as previous studies have reported (Engl et al., 2013; Nico Papinutto et al., 2015, 2020; Solstrand Dahlberg et al., 2020). The highest correlation was found with thalamus volume (*Pearson’s r* = 0.51). No significant correlation was found with ventricular CSF volume. Correlations with height and weight are low.

We found a statistical difference between SC CSA between male and female; females have a significantly smaller CSA than males as previous studies have shown (Nico Papinutto et al., 2015, 2020; Solstrand Dahlberg et al., 2020).

Regarding the effect of age on SC CSA, we found a decrease of CSA with age. Linear and quadratic fit gave very similar results (*R*^*2*^ = 0.031 for both fits). Since the age range of the participants goes from 48 to 80 years old, it is not surprising that a linear fit is also adequate, in comparison with results reported by others (Kesenheimer et al., 2021; Nico Papinutto et al., 2020). Since CSA peaks around 45 years old (Kesenheimer et al., 2021; Nico Papinutto et al., 2020), CSA values for the age range of this study are decreasing with age as we observed (see **Figure 5**).

### 4.4. Normalization methods

Stepwise linear regression led to thalamus volume and brain volume as predictors for SC CSA. We show for the first time thalamus volume as a predictor for normalization of SC CSA. This model significantly reduced inter-subject variability; COV went down from 10.09% to 8.64 %, which represents a reduction of 14.37%. Other parameters were not significant since they were excluded during the stepwise model (*p-value* > 0.05). Sex alone was not a significant predictor (*p-value* > 0.05) even if there is a significant difference between male and female CSA. Note the strong correlation between sex and thalamus volume (*Pearson’s r* = 0.36) and between sex and brain volume (*Pearson’s r* = 0.49). When adding the interaction bewteen sex and brain volume, sex and the interaction became significant. The effect of brain volume on SC CSA varies between male and female as we can observe in **Figure 6**. The interaction between thalamus volume and sex wasn’t significant. The model including brain volume, thalamus volume, sex and sex/brain volume interaction led to a COV of 8.59%, a reduction of 14.85 %. Including sex and brain volume interaction led to the best COV reduction. To our best knowledge, interaction of sex and brain volume was never considered in previous normalization models, only the fixed effect of sex on CSA was included. These findings reveal the importance to consider that factors can vary differently for males and females. We also proposed a model without the thalamus volume, given the difficulty to measure it (it requires the proper anatomical sequence with sufficient contrast and resolution). This model reduced CSA variability less than when including thalamus volume (11.22% of reduction). The combination of thalamus volume, brain volume and sex better explains CSA variability.

We obtained a lower reduction than other models presented in previous studies (Kesenheimer et al., 2021; Nico Papinutto et al., 2020). Kesenheimer *et al*. (2021) obtained a reduction of COV of 23.7% using sex, brain WM volume and SC canal area, Papinutto *et al*. (2020) obtained a reduction of 17.74% using V-scale and axial-canal product. It is important to consider that the predictors of the normalization methods were different, mainly regarding metrics related to the SC canal which could explain the smaller reduction of COV obtained with our model (14.85%). We did not include SC canal metrics in our analysis because of the lack of automated methods for robust SC canal segmentation combined with the large number of participants in this study. Also, the size of the cohort is larger than in the previously mentioned works, N=60 for (Kesenheimer et al., 2021) and N=129 for ((Nico Papinutto et al., 2020), which impacts the distribution and coverage of the data of the participants and affects the normalization method.

Age was not a significant predictor for SC CSA. Trends for CSA and brain volume for the age range of our study are very similar. The effect may differ for younger people since brain volume decreases linearly with age while CSA increases until about 45 years old and decreases afterward (Nico Papinutto et al., 2020).

We have to consider the fact that older people may be more subject to motion in the MRI than younger people (discomfort, difficulty breathing) resulting in a bias in the measured CSA (blurring, motion artifact ghosting). Further investigations are needed to validate if the model can expand to ages not included in this study.

The normalization model was generated from T1w data with a specific protocol. Subsequent studies should assess whether the model is adequate for other acquisition parameters and contrasts. It is known that the output CSA varies for different acquisition protocols (Cohen-Adad et al., 2021). However, since there is a direct relationship between CSA values from different contrasts, there would be a systematic offset in the produced CSA. Since the model is linear, it should hold for different contrasts.

The model was developed from healthy participants; the question remains if it would be applicable to patients with neurodegenerative diseases such as MS. Brain volume changes have been associated with atrophy for various neurodegenerative diseases. A normalization model including brain volume may not be generalizable for those patients. As done in other studies (Kesenheimer et al., 2021), intracranial volume could be a relevant substitute for brain volume since it is not affected by neurodegenerative diseases. Including sex interaction here will also be important and can improve the normalization model. Further studies could include intracranial volume in the normalization model to make it available in the software SCT.

Furthermore, other confounding factors could possibly affect image acquisitions for pathological patients. Severe motor disability could induce some breathing difficulties which induces considerables motion artifacts, thus SC segmentation and CSA bias.

### 4.5. SCT normalization feature

We made available the obtained normalization model in the open-source software SCT within `sct_process_segmentation`. Since thalamus volume may not be available in all SC MRI studies, we made it possible for the user to normalize CSA values without thalamus volume. Even if the best model was obtained with thalamus volume, brain volume, sex and sex interaction between brain volume and sex, normalizing SC CSA without thalamus volume could still reduce CSA variability (reduction of 11.22%). The normalization feature can be used by adding the option `-normalize` followed by the predictors and their corresponding values. For more information on usage, refer to: https://spinalcordtoolbox.com/user_section/command-line.html#sct-process-segmentation.

## 5. Conclusions

This study features an analysis of factors contributing to SC CSA variability at a larger scale than what was done previously to our best knowledge. We introduced a new reference for CSA measurements based on a neurological reference (PMJ) to overcome vertebral reference limitations (neck flexion and extension). We computed over a large cohort of participants SC CSA at 64 mm from the PMJ on T1w scans from the UK Biobank database. No significant age trend was found while SC CSA was significantly different for males and females. We present an effective normalization model including thalamus volume, brain volume, sex and sex/brain volume interaction readily usable in SCT. The most relevant factors to explain SC CSA variability are related to the brain; these findings show the importance of having a brain MRI acquisition in SC studies/research. Reducing inter-subject variability could improve comparison between CSA measures to increase its sensitivity and specificity to better assess pathology-related changes.

## Abbreviations

BMI: Body Mass Index
COV: Coefficient of Variation
CSA: Cross-Sectional Area
CSF: Cerebrospinal Fluid
GM: Gray Matter
MS: Multiple Sclerosis
PMJ: Pontomedullary Junction
SC: Spinal Cord
SCT: Spinal Cord Toolbox
WM: White Matter

## 6. Acknowledgements

We thank Étienne Bergeron, Olivier Lupien-Morin, Benjamin Carrier and Yassine El Bouchaibi for helping with the manual correction and labeling, Nicholas Guenther and Alexandru Foias for data management with git-annex. This research was conducted using data from UK Biobank, a major biomedical database (www.ukbiobank.ac.uk). This study was funded by the Canada Research Chair in Quantitative Magnetic Resonance Imaging [950-230815], the Canadian Institute of Health Research [CIHR FDN-143263], the Canada Foundation for Innovation [32454, 34824], the Fonds de Recherche du Québec - Santé [28826], the Natural Sciences and Engineering Research Council of Canada [RGPIN-2019-07244], the Canada First Research Excellence Fund (IVADO and TransMedTech), the Courtois NeuroMod project, the Quebec BioImaging Network [5886, 35450], the Spinal Research and Wings for Life (INSPIRED project).

## 8. Supplementary material

**Figure S1.**
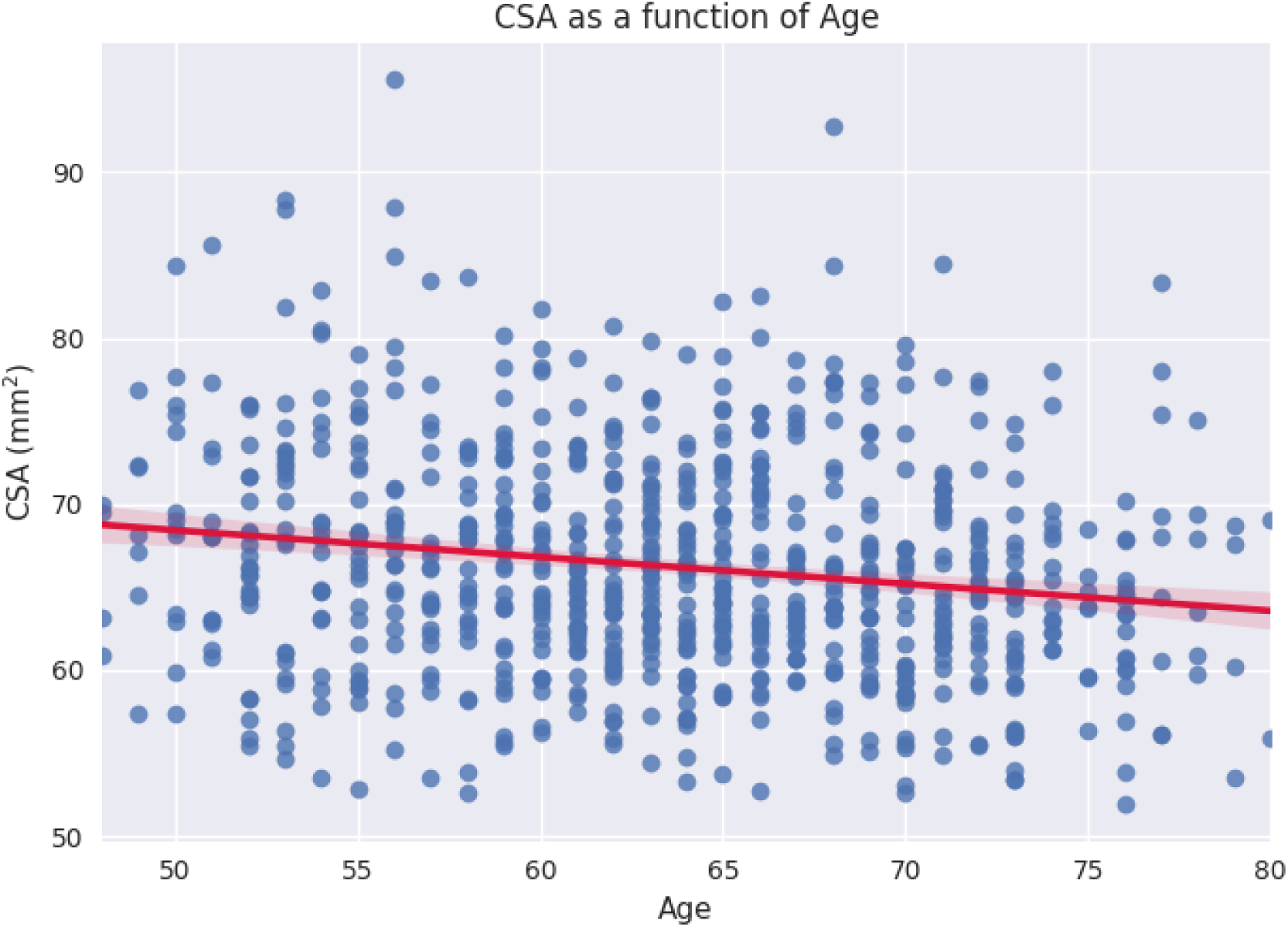
Scatterplot of CSA as a function of age.

**Figure S2.**
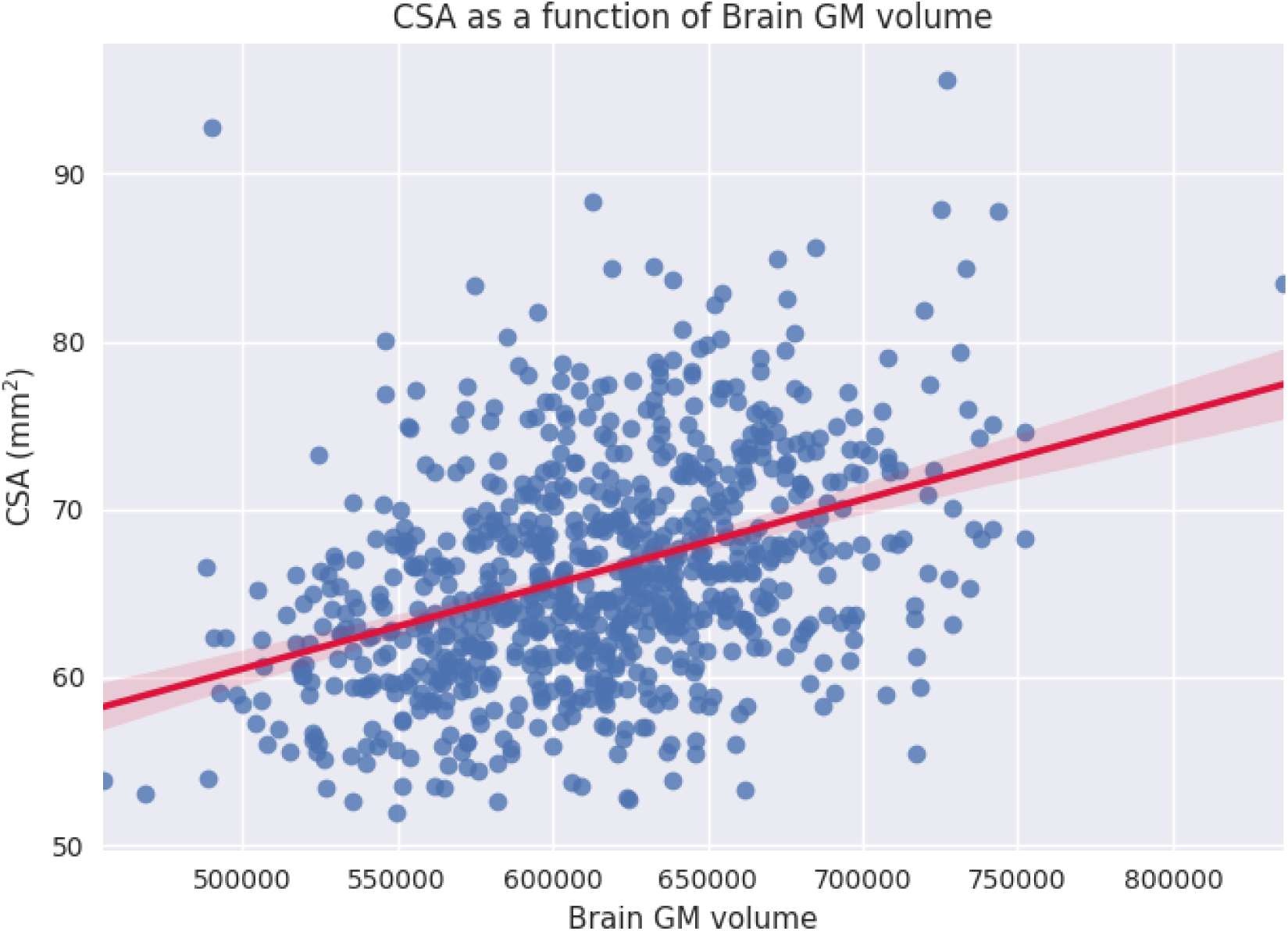
Scatterplot of CSA as a function of brain GM volume.

**Figure S3.**
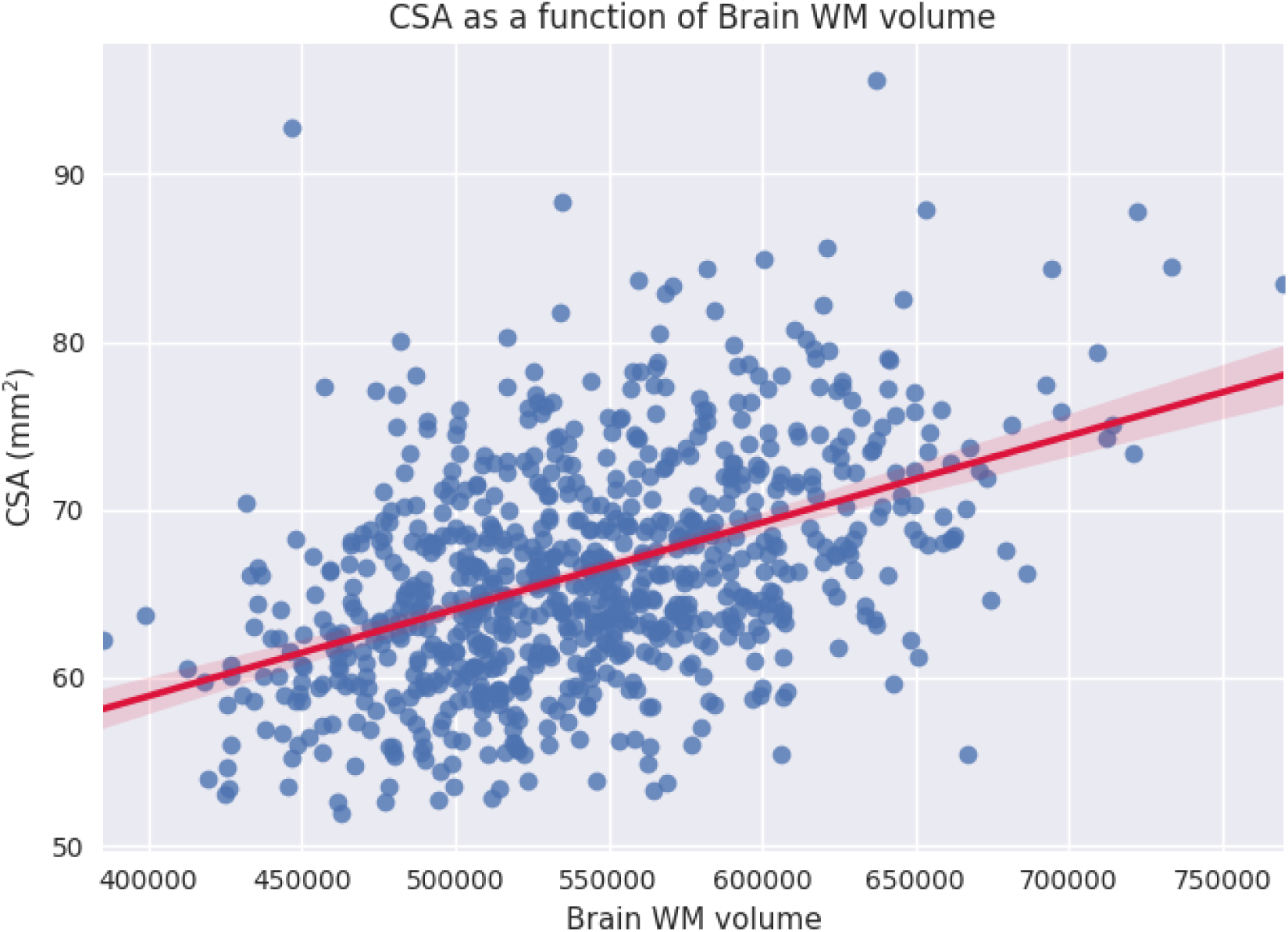
Scatterplot of CSA as a function of brain volume WM volume.

**Figure S4.**
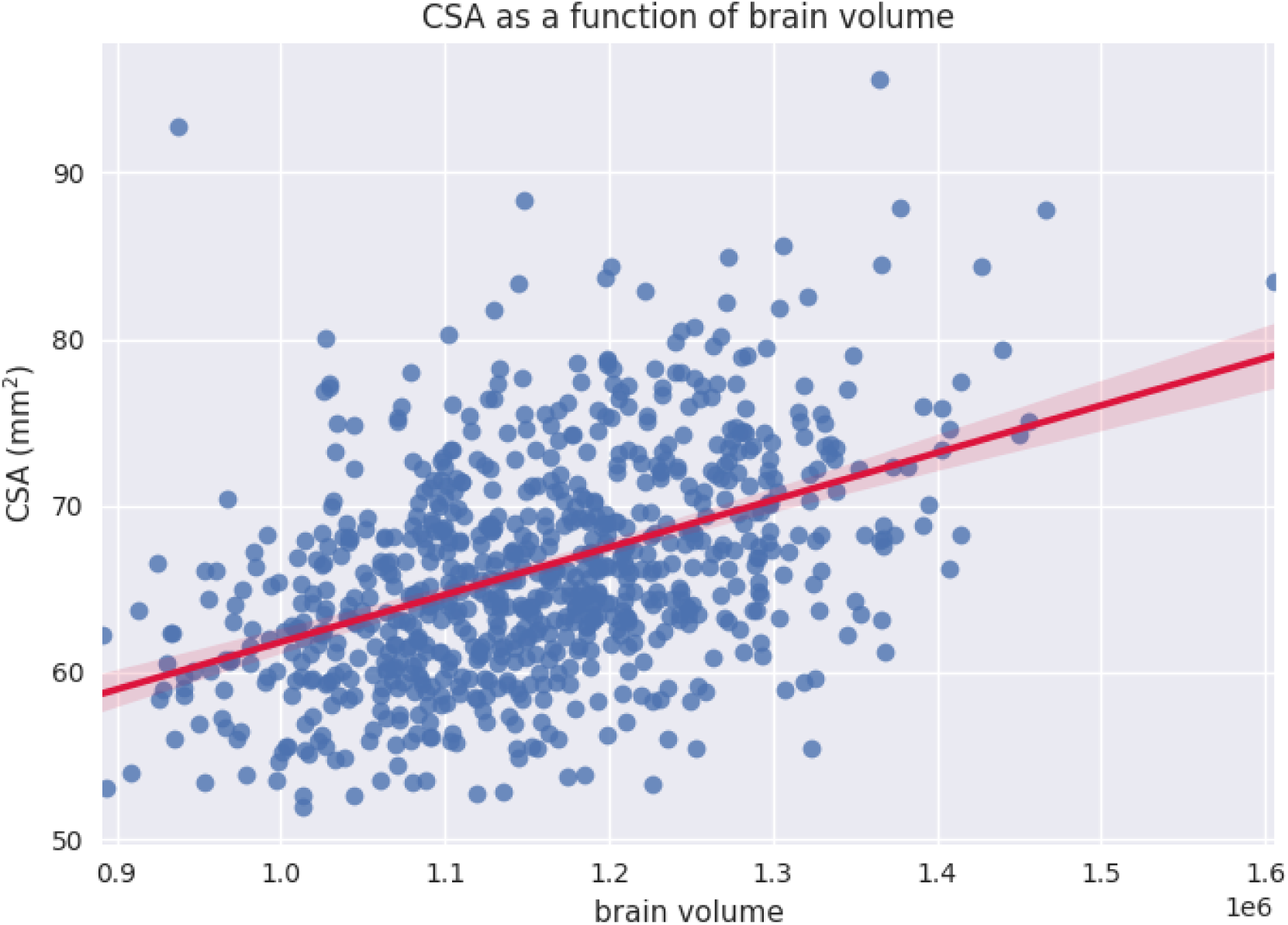
Scatterplot of CSA as a function of brain volume.

**Figure S5.**
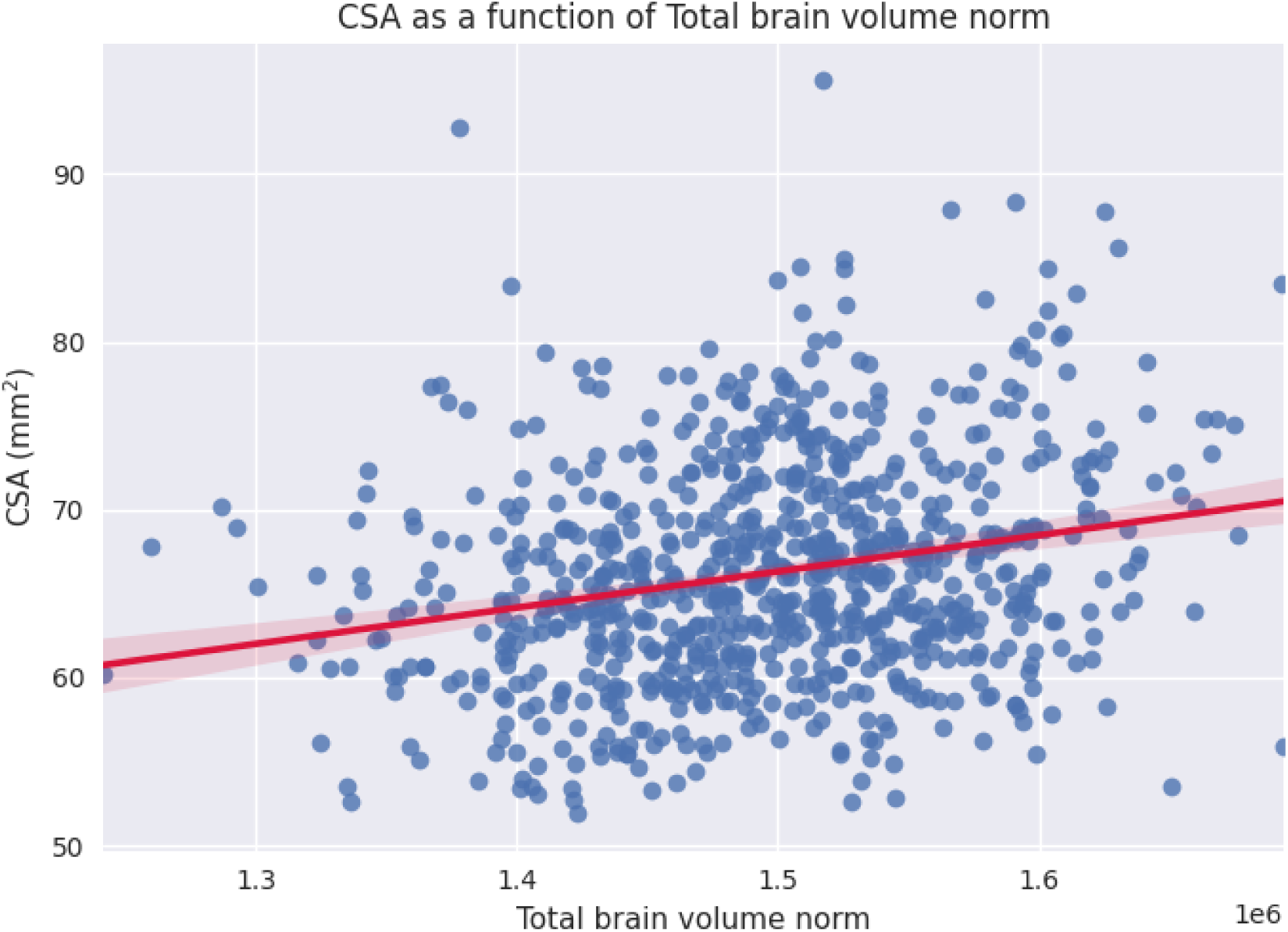
Scatterplot of CSA as a function of brain volume normalized for head size.

**Figure S6.**
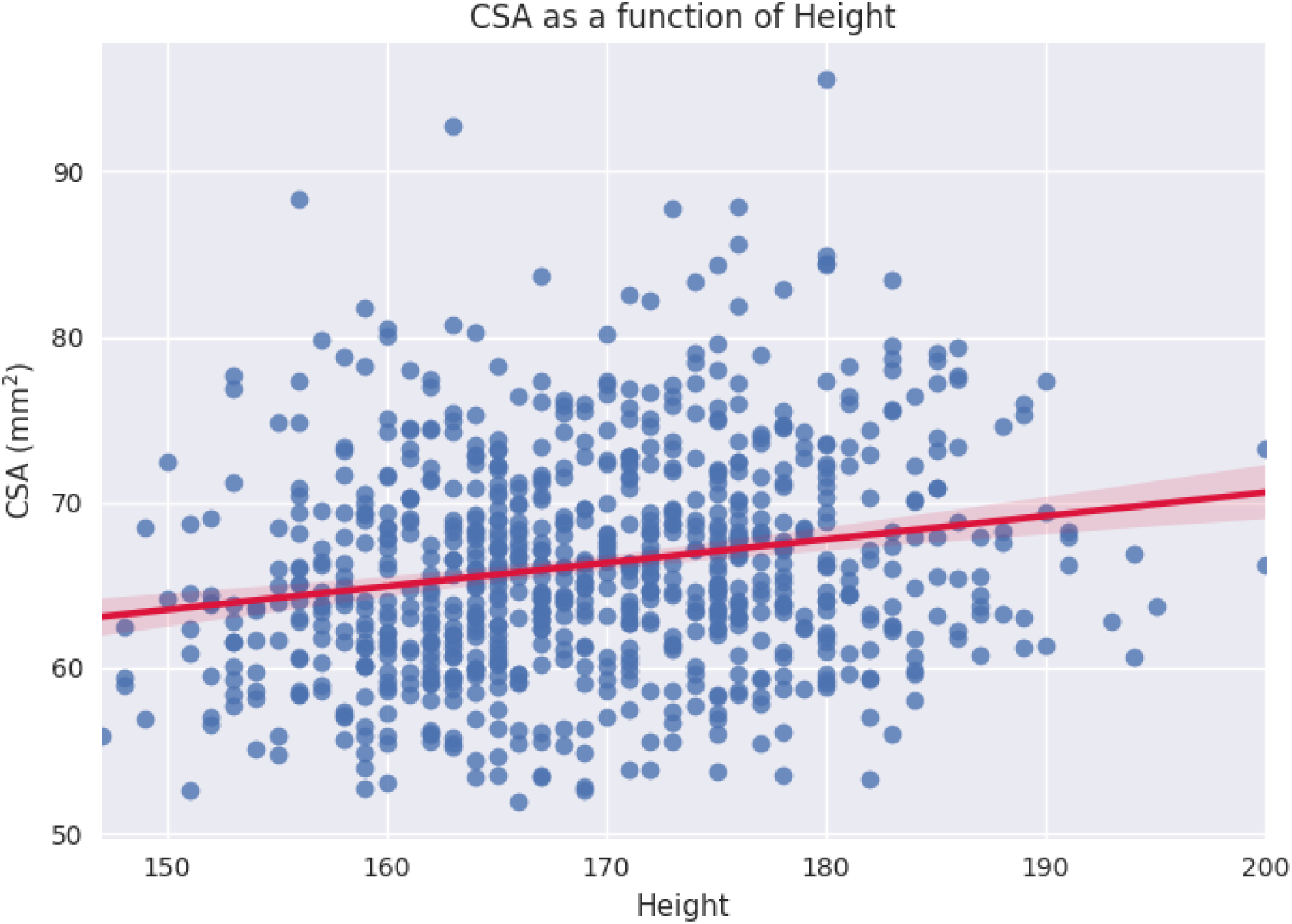
Scatterplot of CSA as a function of height.

**Figure S7.**
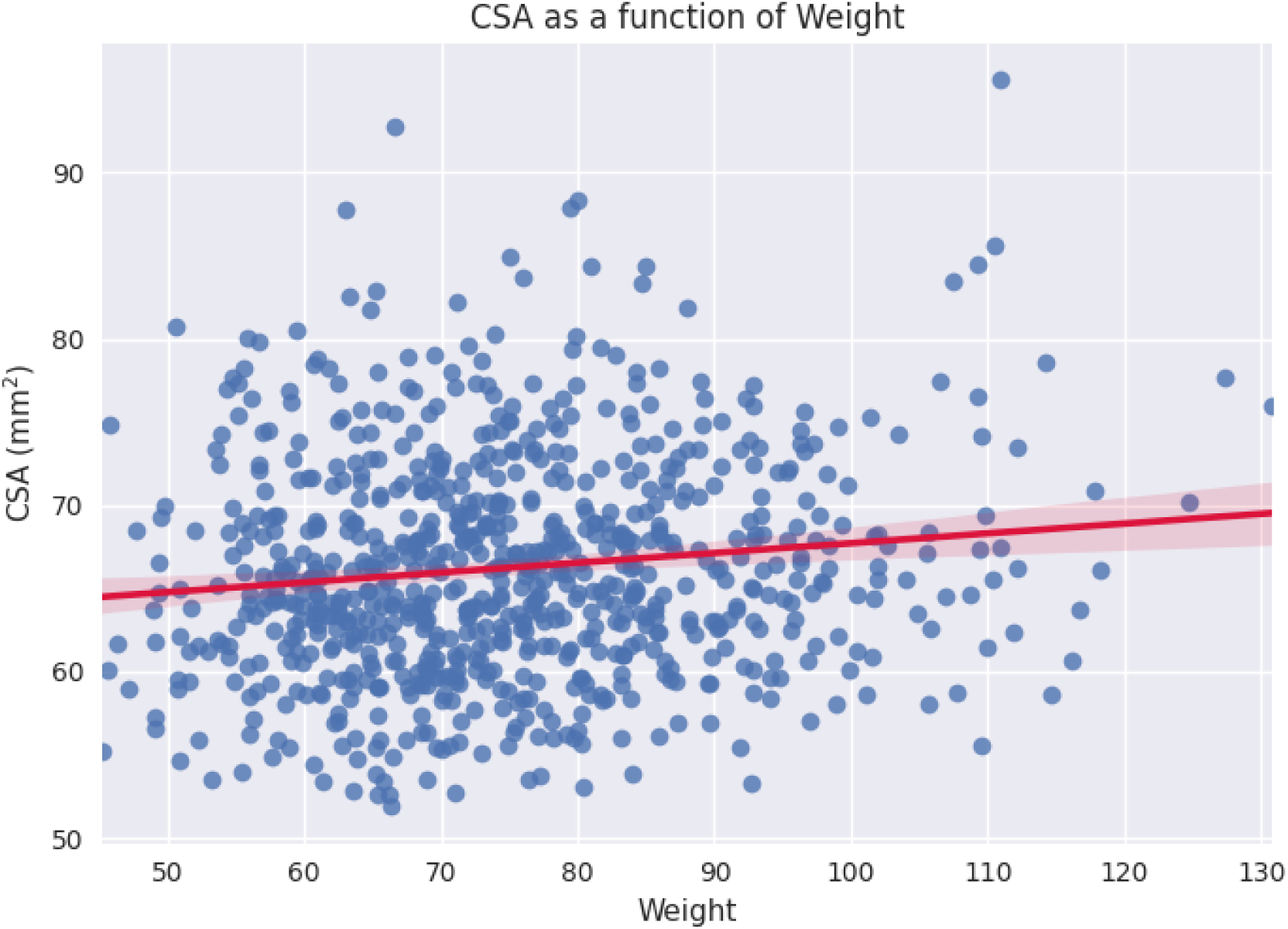
Scatterplot of CSA as a function of weight.

**Figure S8.**
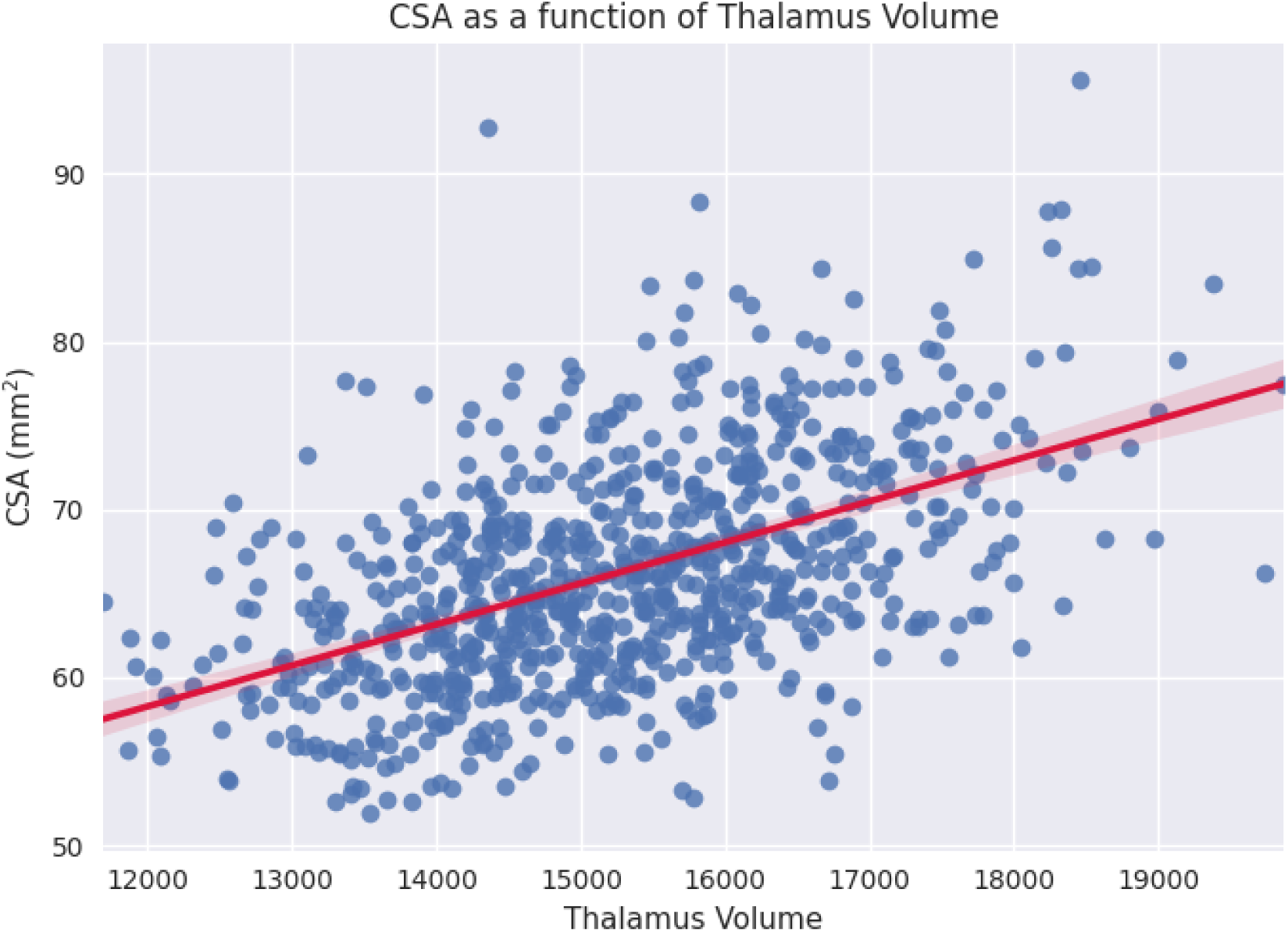
Scatterplot of CSA as a function of thalamus volume.

**Figure S9.**
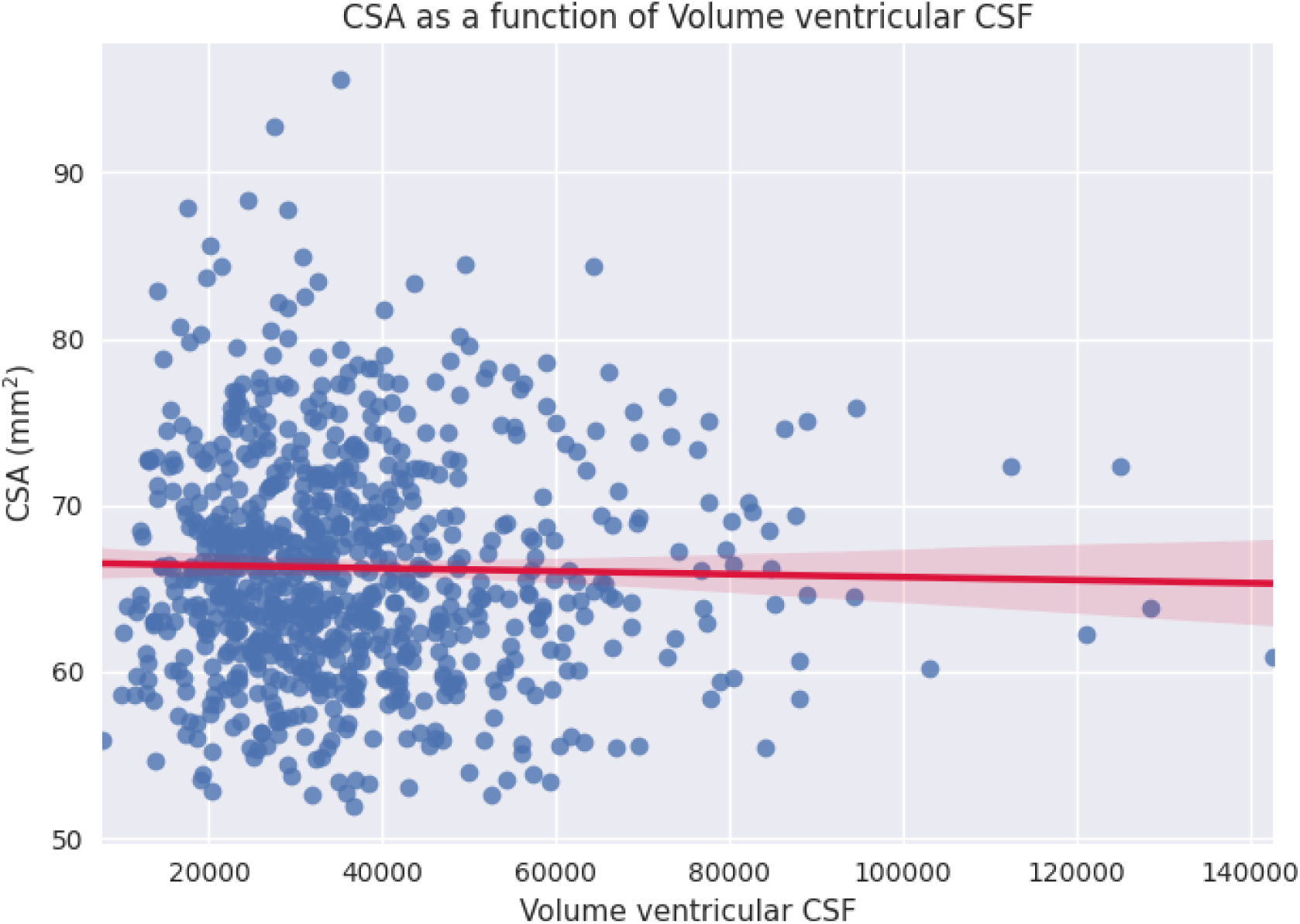
Scatterplot of CSA as a function of ventricular CSF volume.

**Figure S10.**
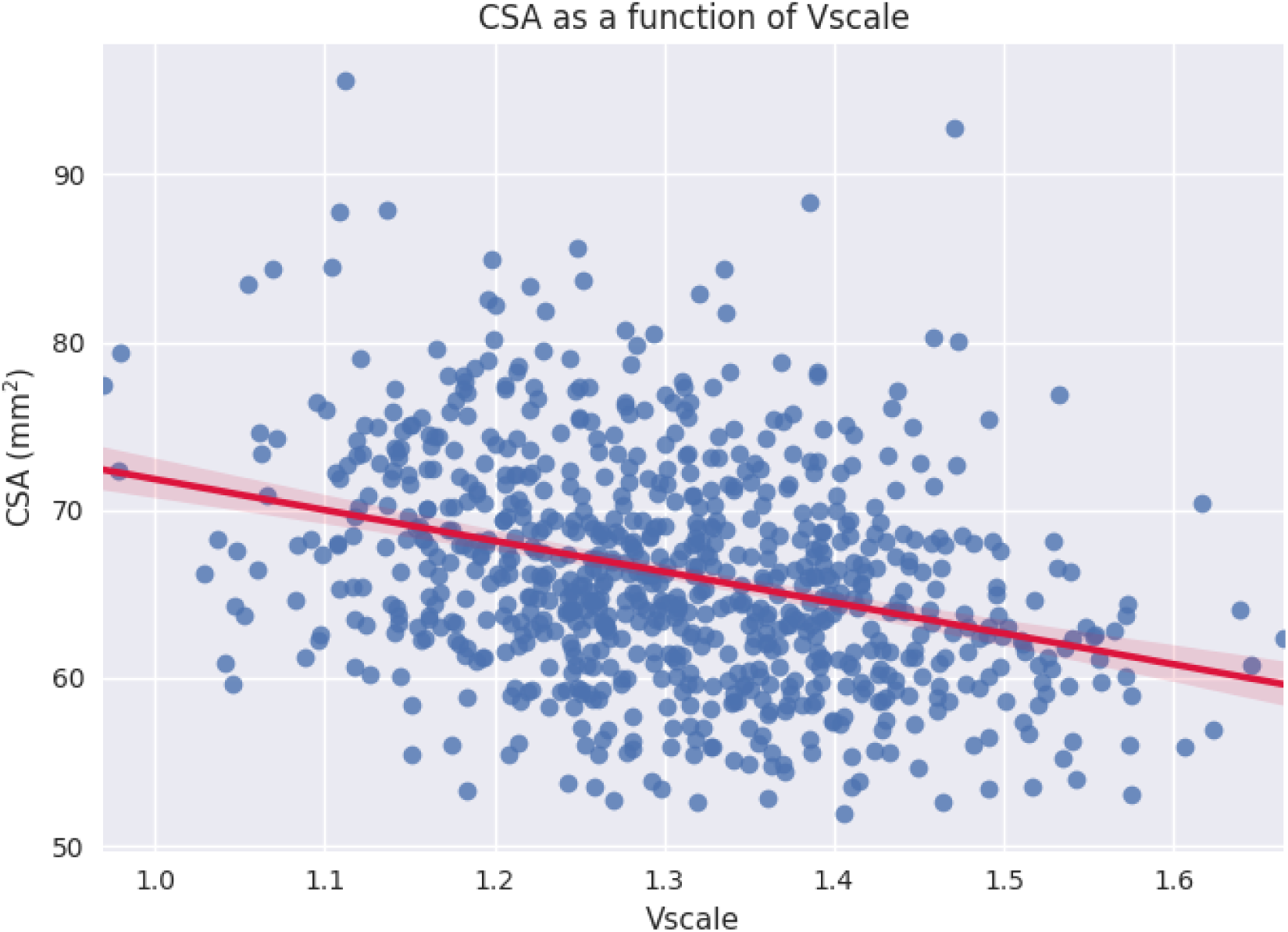
Scatterplot of CSA as a function of v-scale.

## Footnotes

1 https://biobank.ndph.ox.ac.uk/showcase/label.cgi?id=2406

2 https://github.com/spine-generic/spine-generic/releases/tag/v2.6

3 https://github.com/neuropoly/spinalcordtoolbox/releases/tag/5.4

4 https://github.com/sct-pipeline/ukbiobank-spinalcord-csa/blob/master/preprocess_data.sh

5 https://github.com/sct-pipeline/ukbiobank-spinalcord-csa/blob/master/process_data.sh

6 https://github.com/sct-pipeline/ukbiobank-spinalcord-csa/blob/master/README.md

7 https://github.com/sct-pipeline/ukbiobank-spinalcord-csa/blob/master/README.md#quality-control

8 https://spine-generic.readthedocs.io/en/latest/analysis-pipeline.html#quality-control

